# Spatiotemporal microhabitat heterogeneity and dispersion patterns of two small mammals in a temperate forest

**DOI:** 10.1101/278390

**Authors:** Ivan Mijail De-la-Cruz, Alondra Castro-Campillo, Alejandro Zavala-Hurtado, Arturo Salame-Méndez, José Ramírez-Pulido

## Abstract

One basic issue in ecology is to understand how seasonal spatiotemporal shifts in habitat heterogeneity shape spatial patterns in the distribution and coexistence of organisms. For example, how do spatial dispersion of syntopic species reflects their respective resource preferences and partition, during microhabitat seasonal shifts? To address this, we assessed changes in microhabitat structure heterogeneity, between the two annual seasons (dry and rainy seasons) in a midlatitude temperate forest by analyzing 23 habitat variables using multivariate statistics. Furthermore, to determine how such microhabitat changes affected the spatial segregation of two congeneric species of small mammals (*Peromyscus difficilis*, *P. melanotis*), we used second-order spatial statistics to detect changes in their microdistribution changes and general linear models to evaluate their habitat preferences. In accordance with their respective morphology and ambulatory habits, the medium-sized, semi-scansorial *P. difficilis* was resident year-round and preferred microhabitats with high amounts of log ground cover, while the small, opportunistic, cursorial *P. melanotis* varied its occupancy area depending on the density of herbaceous and cover of woody plants. Both species showed more similar microdistribution patterns under the more benign microhabitat conditions of the rainy season (denser plant coverage, humid-cool weather), but became more segregated form each other during the less favorable conditions of the dry season (scarcer plant coverage, dry-cold weather). Therefore, seasonal changes in microhabitat heterogeneity influenced the dispersion patterns of both *Peromyscus* species and revealed their temporal preferences for microhabitat resources.

## Abstract

**Resumen.** Un tema básico de la ecología es comprender cómo los cambios espacio-temporales de la heterogeneidad del microhábitat, dan forma a los patrones espaciales de distribución y coexistencia de los organismos. En otras palabras, cómo es que la dispersión espacial de especies sintópicas refleja sus preferencias y partición de recursos durante los cambios estacionales del microhábitat? Para abordar esto, evaluamos los cambios en la estructura vertical-horizontal del microhábitat (heterogeneidad) a lo largo de las dos estaciones anuales (temporadas seca y lluviosa) de un bosque templado, analizando 23 variables con estadística multivariada. Además, para determinar cómo tales cambios del microhábitat afectaban la segregación espacial de dos especies congenéricas de pequeños mamíferos (*Peromyscus difficilis*, *P. melanotis*), utilizamos estadística espacial de segundo orden y modelos lineales generalizados para detectar sus cambios específicos de microdistribución y sus preferencias de hábitat, respectivamente. De acuerdo con su morfología y hábitos ambulatorios respectivos, el mediano y semiescansorial *P. difficilis* fue residente durante todo el año y prefirió los microhábitats con una alta cobertura de troncos sobre el suelo, mientras que el oportunista, pequeño y cursorial *P. melanotis*, varió su área de ocupación, dependiendo de la densidad de plantas herbáceas y la cobertura de plantas leñosas. Bajo condiciones de microhábitats más benignas en la época de lluvias (mayor densidad de cobertura vegetal, clima húmedo-fresco), ambas especies mostraron patrones de microdistribución más próximos, pero se volvieron más segregadas entre sí durante las condiciones menos favorables de la temporada seca (cobertura vegetal más escasa, clima seco-frío). Por lo tanto, los cambios estacionales en la heterogeneidad del microhábitat influyeron en los patrones de dispersión de ambas especies de *Peromyscus* y revelaron sus preferencias temporales por los recursos de microhábitat.

## Introduction

Spatiotemporal habitat heterogeneity has a substantial impact on species coexistence. For example, species may become spatially segregated according to niche preferences, such as resource requirements (Valladares *et al*. 2015). Model simulations reveal the potentially important role of spatial heterogeneity and its complex and delicate interplay with dispersal in mediating long-term outcomes of species coexistence (Schreiber and Killingback 2013; Valladares *et al*. 2015). For instance, an increase in the number of habitat types, resources, and structural complexity should increase available niche spaces, thus allowing more species to coexist (Currie 1991). Equally crucial for the maintenance of species coexistence is habitat heterogeneity over time, whose influence on natural communities also varies depending on the temporal scale (Valladares *et al*. 2015). Temporal fluctuations in habitat structure can stabilize coexistence via the “storage effect” (Chesson 2000), when inter-annual variation in climate or resource availability favors one group of species over others (Zavaleta *et al*. 2003).

Heterogeneous habitats contain patches of varying size, distribution, resources, environmental conditions, and species composition (Fahrig and Merriam 1994) that vary spatiotemporally, depending on observation scale and habitat type (Wiens 2000). It is highly likely that small body size species such as small mice perceive spatial habitat heterogeneity at a fine scale (microhabitat). Therefore, both movement and foraging by this kind of small mammals are affected by habitat heterogeneity (Bowne *et al*. 1999), having different responses among species or demographic groups within them (Dooley and Bowers 1998). Here we focus on whether small mammals are capable of perceiving and responding to spatiotemporal microhabitat heterogeneity.

Small mammals use resources selectively, based on their requirements for growth, survival, and reproduction (Johnson and Gaines 1980). For instance, spatiotemporal habitat heterogeneity affects *Peromyscus* mice in terms of species coexistence (M’Closkey and Fieldwick 1975; M’Closkey and Lajoie 1975), spatial segregation (Monamy and Fox 1999), and competition (Morris 1984a; Seagle 1985). Indeed, several studies have quantified variation in resource use in heterogeneous environments within a community among mice from the same family or genus (Kaufman and Kaufman 1989 and references therein). Indeed, closely related species are especially valuable for studies of spatial and resource partitioning, since they are most likely to be current or past competitors (Kalcounis-Rüppell and Millar 2002). Therefore, since habitat use varies spatiotemporally (Haim and Rozenfeld 1993) due to distribution and availability of resources, we expect this variation to affect the use of resources and relationships among closely related species.

In this study, the primary objective was to analyze how microhabitat heterogeneity is structured between the dry and rainy seasons of a temperate mixed forest, and how these changes affect the coexistence of *Peromyscus difficilis* and *P. melanotis*. To do so, we addressed four main aspects; first, we assessed the presence of spatiotemporal heterogeneity at a fine scale (microhabitat) according to vertical and horizontal structure indicators. Second, we determined the ecological dispersion patterns of *P. difficilis* and *P. melanotis* and tested whether the use of space by these two *Peromyscus* was affected by spatiotemporal changes in microhabitat heterogeneity. Third, we determined what kind of spatial relationships occurred between the two species depending on seasonal changes in microhabitat heterogeneity (*i. e*., attraction/positive: sharing microhabitats and resources, or repulsion/negative: not sharing). Finally, we assessed what particular structural elements of each one of the microhabitats best explained capture frequency for each *Peromyscus* species in each season.

## Materials and methods

### Study area

The study area was located in a temperate mixed forest (coniferous and broad-leaved trees) at Desierto de Los Leones National Park (DLNP, CONANP 2006), Mexico City, within the Transmexican Neovolcanic Range (Universal Transverse Mercator, UTM, boundary coordinates: 465261.25 m E, 2137029.52 m N; 468996.54 m E, 2129839.47 m N). The isothermal temperate climate belongs to the most humid of the subhumid type with summer rainfalls (C(W2) W (b)ig); winter rainfall accounts for < 5% of total annual rainfall (CONANP 2006). There is an annual accumulated precipitation of 1340.6 mm (monthly mean 111.72 ± 109.1; minimum 10.3 in December, maximum 275.9 in August). The rainy season (CONANP 2006) is from summer through early fall (June - September) with humid-cool weather and a monthly average precipitation of 252.92 ± 28.01 (218.8, September - 275.9, August) and an average monthly temperature (°C) of 11.72 ± 0.53 (11.3, September - 12.5, June). The dry season (CONANP 2006) occurs from late fall through winter (November - February) with cold, dry weather: monthly average precipitation (mm) from November - January is 13.2 ± 3.11 (10.3, December - 16.5, November), and average monthly temperature (°C) is 8.97 ± 0.68 (8.3, January - 9.7, November). The dominant vegetation in most of DLNP is *Abies-Pinus-Quercus* forest, at 2,800 - 3,000 m (CONANP 2006). According to official sources (CONANP 2006) and our unpublished data, the most conspicuous plant species by stratum are as follows. The canopy includes Christmas fir (*Abies religiosa*), several species of pine trees, including *Pinus patula, P. hartwegii, P. leiophylla,* and *P. montezumae*, and oak trees *Quercus laurina, Q. castanea, Q. laeta*, together with other broad-leaved trees (*e. g., Prunus serotina, Salix paradoxa*, and *Buddleia cordata*). Among understory shrubs are *Senecio barba-johannis, Symphoricarpos microphyllus, Cestrum anagyris, Solanum cervantesii, Physalis viscosa, Fuchsia microphylla*, *Baccharis conferta*; there are also isolated elements of *Berberis moranensis, Garrya laurifolia, Arbutus xalapensis, Buddleia cordata*, and *Clethra mexicana* within the oak trees. Among the most frequent herbs are *Acaena elongata, Sigesbeckia jorullensis, Alchemilla procumbens*, *Geranium seemannii, Valeriana clematitis*, and *Archibaccharis hirtella*. The ground level bears a rich variety of mosses (*e. g., Aloinella catenula, Anacolia laevisphaera, Bartramia* spp., *Bryum procerum, Grimmia* spp., *Herzogiella cylindricarpa, Mironia* spp.) and fungi, whose fruiting bodies are conspicuous during the rainy season (*e. g*., *Cantharellus cibarius, Phlogiotis helvelloides, Amanita* ccf. *citrina, Geastrum triplex, Ramaria* sp.).

### Microhabitat features

We set a 2,475 m^2^ (55 x 45 m) surface plot at 2,289 m (UTM 2144958 m N, 14477883 m E) (Figure 1). The plot was gridded with 12 vertical lines (A - L) by 10 horizontal lines (1 - 10), with lines placed every 5 m. Intersections between vertical/horizontal lines were marked with buried wooden stakes (150 x 2.5 x 2.5 cm) to construct a coordinate system for 120 independent sampling stations (Figure 1). To assess spatiotemporal heterogeneity of microhabitats within the entire plot, we delimited an influence zone (Figure 1), drawing a 2.5 m^2^ square around each sampling station. We used these influence zones to sample microhabitat characteristics around each sampling station, using 23 environmental variables (Table 1) for ten days: once in the dry season (July) and once during the rainy season (February; CONANP 2006). All selected variables qualify as components of the vertical and horizontal structure of microhabitats, as well as factors affecting rodent distribution at this spatial scale (Morris 1984b, 1987; Jorgensen 2004; Coppeto *et al*. 2006; Villanueva-Hernández *et al*. 2017). Further, taken together, these 23 variables (Table 1) provide information about possible refuges for the small mammals from predators, as well as spaces for resting, breeding, food resources, and safe routes which can be used to move from one place to another to avoid predation risk (Jorgensen 2004). More detailed information about the ecological significance of each variable appears in Table 1.

**Figure 1.**
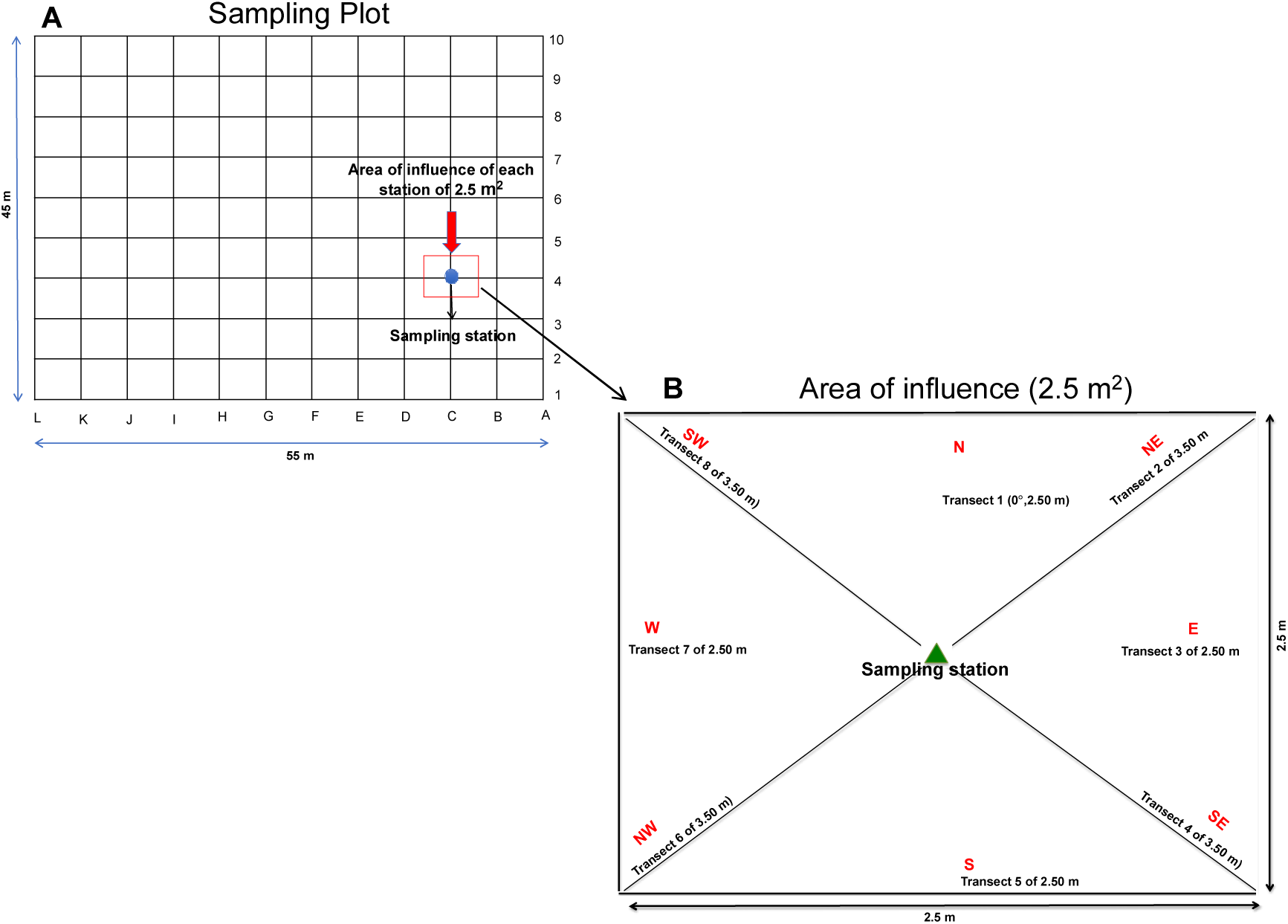
Sampling plot design (A). Total plot area was 2475 m^2^; each intersection (dots, n = 120) was an independent sampling station for capturing mice. Influence area of 2.5 m^2^ around each station sampling (central green triangle) (B), where 23 environmental variables of microhabitat were sampled along eight transects (see text).

**Table 1.**
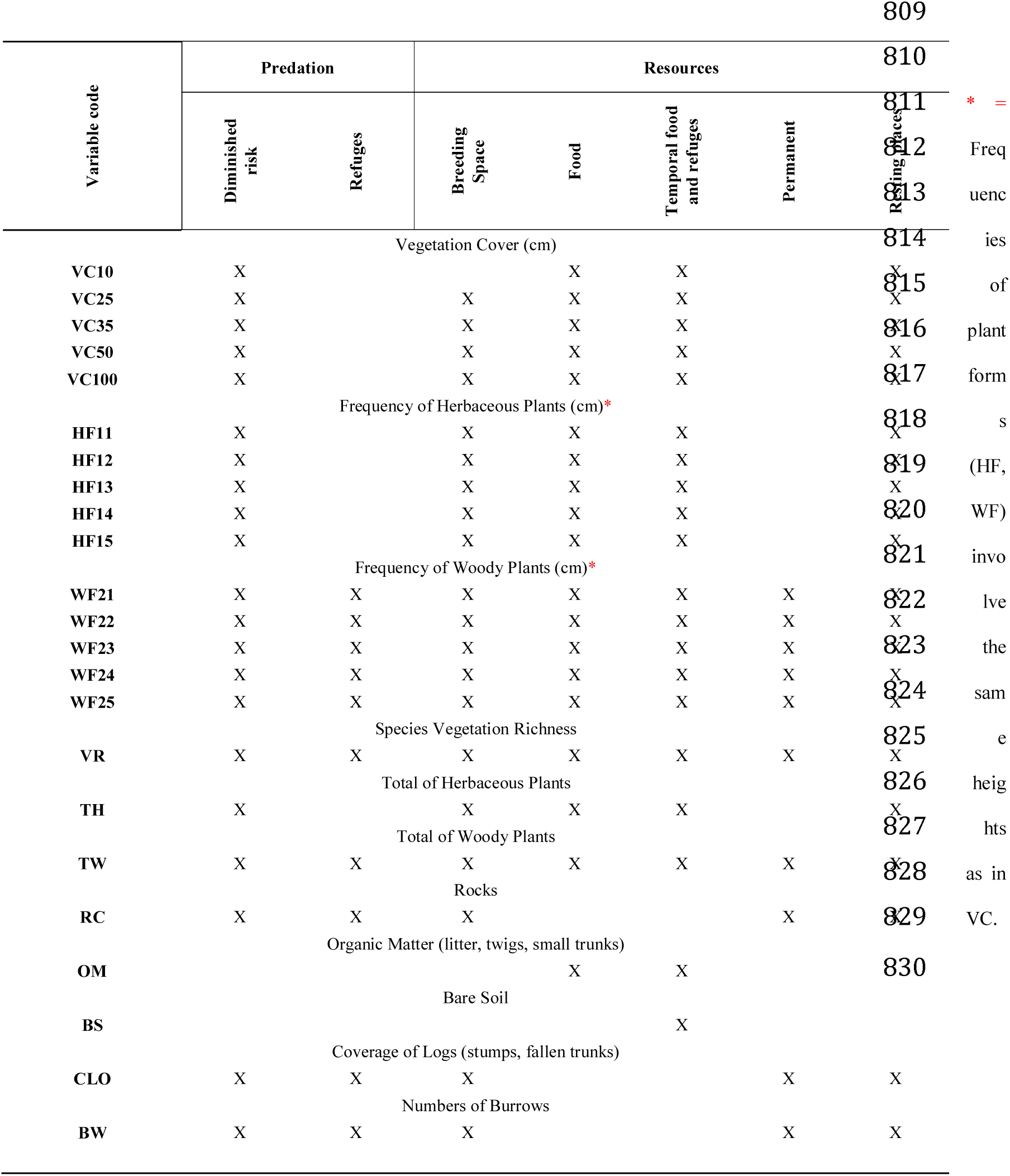
Environmental variables used to measure the horizontal and vertical structure of microhabitat at each sampling station, showing its ecological interpretation (kind or resource type) for both species of *Peromyscus*.

The variables we sampled include the following: Vegetation Cover at five different heights at the understory in cm: *i. e*., 10, 25, 35, 50, and 100 cm (VC10 - VC100; *e. g*., VC10 is vegetation coverage at 10 cm height). Number of herbaceous forms at five different heights: *i. e*., 10, 25, 35, 50, and 100 cm (HF1-5; *e. g*., HF3 is number of herbaceous plants at level 3, or 25 cm height). Number of woody forms (WF1-5; *e. g.,* WF5 is the number of woody forms at level 5, or 100 cm height). Total Herbaceous plants (TH) and Total Woody plants (TW) are the total number of each type counted within each influence zone. Vegetation species richness (VR) is total number of plant species. Other variables included measurements of the coverage of fixed microhabitat elements (rocks = RC, logs = CLO, stumps and fallen branches), organic matter (OM, mostly leaf litter and small twigs), and bare soil (BS). Finally, we counted the number of mouse burrows (BW), recording each hole in the ground *ca*. 8 - 12 cm of diameter within the influence zone (Álvarez-Castañeda 2005; Fernandez *et al*. 2010).

To measure all variables for each season in all influence zones, we adapted Canfield’s (1941) Line Intercept (LI) method, which allowed us to sample the within-plot variation and quantify microhabitat changes in plant species cover and height over time. We drew eight graduated transects (cm) within the respective influence zone of each station to cover the entire surface area (Figure 1): four 2.5 - m transects were cardinally oriented (N, S, E, W), while the other four (3.30 m long) were diagonally oriented (NE, NW, SE, SW). We measured plant coverage at the five different heights (VC10 - 100), as well as coverage of fixed elements; *i. e.,* rocks (RC), logs (CLO), bare soil (BS), dead organic matter (OM) for all transects. We only counted plants intercepted by diagonal transects for vegetation species richness (VR), for number of plant life forms at the different height levels (herbaceous, HF1 - 5; woody, WF1 - 5), and for total herbaceous (TH) and woody plants (TW). We calculated the percent coverage of each variable within the influence zones, using the formula 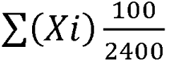, where *Xi* is the length along the transect occupied by each plant in centimeters, and 2,400 is the total length (sum of the eight transects).

### Mouse sampling

We captured *Peromyscus* mice alive over ten months to include data for the dry (October 2013 to February 2014) and rainy (March 2014 to July 2014) seasons. We set a single live trap (H. B. Sherman, Inc., Tallahassee, FL 32303, USA), baited with oat flakes and vanilla extract (Figure 1) at each independent sampling station of the plot (n = 120) (Figure 1). Traps were set for two consecutive nights, for a total of 20 capture events and 2, 400 trap-nights. To record the capture frequency of mice in all the stations for every capture event, we recorded the sampling station coordinates at which mice were caught. We also recorded the species, sex, age, reproductive condition, and any other conspicuous trait that allowed us for individual recognition of every mouse by visual inspection; *e. g.*, acquired traits, such as scars and missing parts in the ears, tail or a missing eye. In addition, to discriminate between new captured individuals and recaptured individuals between capture events, we stained the abdomen of each captured mouse with gentian violet. Since temporary marking with gentian violet lasted in average 75 ± 7 days in the rainy season and up to 180 ± 8 days in the dry season, it proved to be a non-intrusive, affordable, and suitable method for our study objectives. When the gentian violet mark began to fade and a mouse was recaptured, the mark was reapplied. This non-intrusive marking allowed us to avoid overestimating the capture frequency by ensuring we never counted the same individual twice during the same capture event. Moreover, our analyses showed the same dispersion patterns regardless of whether we used both the frequency of newly-captured individuals and recapture frequencies during all capture events, or only new captures in our analyses. In addition, to avoid recurrence behavior (*e. g.,* mice coming back to the traps for bait) or shyness (*e. g.,* mice avoiding traps due to other mice odors), all the traps were thoroughly cleaned and randomly oriented within the influence zone in each capture event. To prevent hypothermia during capture, we placed 3 - 5 cotton balls inside the trap and put the traps inside open plastic bags. Mice capture and handling followed the guidelines of the American Society of Mammalogists (Sikes 2016). Any mice that died were prepared as voucher specimens and incorporated into the Mammal Collection of the Universidad Autónoma Metropolitana-Iztapalapa (UAMI, Ramírez-Pulido *et al*. 1989). A scientific collecting permit, SEMARNAT – 08 – 049 - B, was issued to Alondra Castro-Campillo (ACC) by DGVS, SGPA - 09712/13, SEMARNAT.

### Statistical analysis

We standardized all variables and run statistical tests with an alpha level of 0.05. Examining the Variance Inflation Factor revealed showed no multicollinearity among the 23 habitat variables (*i. e.,* all values were at a threshold < 6, Zar 1999). To identify and categorize microhabitat heterogeneity within the plot grid, we performed two independent Hierarchical Cluster Analysis (HCA) in JMP® (ver. 9, SAS Institute Inc., Cary, NC, 1989 - 2007), one for each season. These analyses enabled us to cluster sampling stations with similar microhabitat characteristics, according to the 23 environmental variables, in each season. We used Ward’s method (1963) to construct the respective dendrograms, where the distance between two clusters is the ANOVA sum of squares between them, added up over all quantitative variables. From visual inspection of each dendrogram and given that there were no drastic changes in the variance scree plot, we obtained three groups of sampling stations for each season, which we interpreted as distinct microhabitat types (M1 - 3). We further statistically validated such microhabitats (Figure A.1) by carrying out independent Discriminant Analyses in each season (DA, Addinsoft SARL’s XLSTAT 2013; F = 9.99 dry; F = 9.64, rains; Wilk’s Lambda = 0.0001 in both). We constructed a different map for each season, assigning each sampling station one of three colors, according to its microhabitat type (M1: red, M2: green, M3: blue, Figure 2). To construct the respective descriptive typologies for the fine scale behavior of each one of the environmental variables, we plotted the 23 standardized variables in the “Y” axis and the microhabitat types in the “X” axis in each season (Figure A.1).

**Figure 2.**
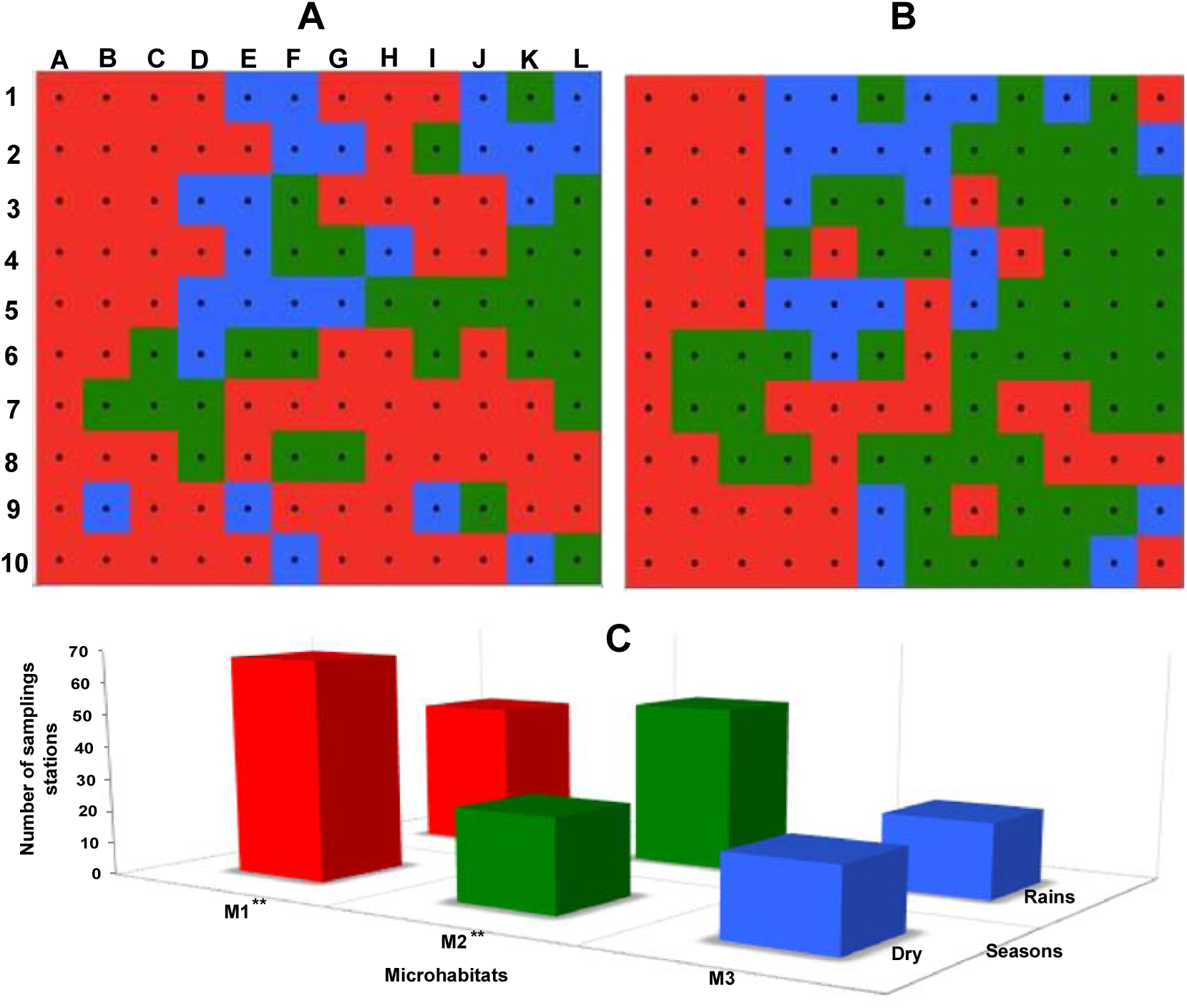
Distribution maps for microhabitats (M1 = red, M2 = green, M3 = blue) within the study quadrant for the dry (A) and rainy (B) seasons. Bar graph showing number shifts of microhabitat types in the dry and rainy seasons (C). Asterisks in M1* and M2* refer to *p* = 0.001.

We calculated descriptive statistics for each variable (*e. g*., mean, median, standard deviation, minimum and maximum values), and tested for normal distribution with Kolmogorov-Smirnov tests (Zar 1999). Then, we analyze distribution changes in each variable between the rainy and dry seasons using Student’s t-test for normally distributed variables and the non-parametric Wilcoxon test for non-normal variables (Table 2). We used a contingency table and *x^2^* test to determine whether the number of stations belonging to each type of microhabitat changed between seasons (seasonal microhabitat heterogeneity). To examine the association between changes in spatial heterogeneity of microhabitats and dispersion patterns of the *Peromyscus* along the entire grid, we conducted a Contingency Table Analysis in each season. In each contingency table, microhabitats were the rows and capture frequency of each *Peromyscus* species were the columns (Table A.2). Then, in order to visualize associations between the *Peromyscus* and microhabitats, we performed a correspondence analysis using the contingency tables for each season (Figure 5). We performed all of these analyses with XLSTAT 13 (Addinsoft SARL).

**Table 2.** Normality (**NOR**) tests for 23 variables (**VAR**) of microhabitat, and comparisons^abc^ (**COM**) for average values between the dry (**DRY**) and rainy (**RAINS**) seasons, respectively. Variable names as in Table 1.

| VAR | NOR<sup>a</sup> |  | COM | VAR | NOR<sup>a</sup> |  | COM | VAR | NOR<sup>a</sup> |  | COM |
| --- | --- | --- | --- | --- | --- | --- | --- | --- | --- | --- | --- |
|  | DRY | RAINS |  |  | DRY | RAINS |  |  | DRY | RAINS |  |
| **BS** | 0.0001 | 0.0001 | *1.0<sup>c</sup>* | **VC100** | 0.002 | 0.003 | 0.0001<sup>c</sup> | **TW** | *0.143* | *0.08* | 0.0004<sup>b</sup> |
| **CLO** | 0.0001 | 0.0001 | *0.972<sup>c</sup>* | **VR** | 0.006 | *0.082* | 0.0001<sup>b</sup> | **WF21** | 0.0001 | 0.0001 | 0.004<sup>c</sup> |
| **OM** | 0.010 | 0.034 | 0.0001<sup>c</sup> | **TH** | 0.0001 | *0.072* | 0.0001<sup>b</sup> | **WF22** | 0.0001 | 0.0001 | 0.026<sup>c</sup> |
| **RC** | 0.0001 | 0.0001 | *0.432<sup>c</sup>* | **HF11** | 0.015 | 0.028 | *0.658<sup>c</sup>* | **WF23** | 0.0001 | 0.0001 | *0.875<sup>a</sup>* |
| **VC10** | 0.002 | 0.041 | 0.0001<sup>c</sup> | **HF12** | 0.0001 | 0.0001 | 0.001<sup>c</sup> | **WF24** | 0.0001 | 0.0001 | *0.378<sup>c</sup>* |
| **VC25** | 0.009 | 0.003 | 0.0001<sup>c</sup> | **HF13** | 0.0001 | 0.0001 | 0.0001<sup>c</sup> | **WF25** | 0.0001 | 0.001 | 0.0001<sup>c</sup> |
| **VC35** | 0.0001 | 0.002 | 0.0001<sup>c</sup> | **HF14** | 0.0001 | 0.0001 | 0.0001<sup>c</sup> | **BW** | 0.0001 | 0.0001 | 1 |
| **VC50** | 0.0001 | 0.019 | 0.0001<sup>c</sup> | **HF15** | 0.0001 | 0.0001 | 0.0001<sup>a</sup> |  |  |  |  |

Parametric: <sup>a</sup>Kolmogorov-Smirnov; <sup>b</sup>Student for two samples; non-parametric: <sup>c</sup>Wilcoxon). The *p*-values in italics do not reject normality (*p* = 0.05), or do not indicate significant differences between pluvial seasons.

### Spatial analysis

To map variations in point density captures of small mammals, and to find density gradients across the plot area, we used the Kernel function (PAST, ver. 3.14, Hammer *et al*. 2001). To test the ecological dispersion patterns of each *Peromyscus* species within the plot (*i. e.,* clustering or overdispersion) in each season, we used a “Nearest Neighbor Analysis” (NNA, Clark and Evans 1954) using PAST software (ver. 3.14, Hammer *et al*. 2001). We applied the wrap-around edge effect adjustment, which is only appropriate for rectangular domains (like our sampling plot; Hammer *et al*. 2001). In general, the NNA compares the mean distance of each focal individual from its nearest conspecific neighbor, with the mean distance expected for a set of points randomly dispersed at the same density (Vázquez and Álvarez-Castañeda 2011). The ratio of the observed mean distance to the expected distance (*R*) indicates how the observed distribution deviates from random; i.e., clustered points give *R < 1*, Poisson patterns give *R ∼ 1*, and overdispersed points give *R > 1*.

To assess whether the two species were associated (attracted) or disassociated (repelled) with each other, we compared dispersion patterns between the two mice, using Ripley’s K bivariate function (Ripley 1976), since the method considers all distances among individuals located under a Cartesian scheme (X, Y) in a quadrat plot (Ripley 1976; Dale 1999; Zavala-Hurtado *et al*. 2000). We used PASSaGE (ver. 2, Rosenberg and Anderson 2011) to carry out Ripley’s bivariate K analysis. We used the option to test the associations conditional on the current locations hypothesis in PASSaGE, where the point locations (sampling stations) remain fixed, and only the point types (each *Peromyscus* species as type points A or B) are randomized; once the number of points of a determined type remains fixed (*Peromyscus* species A), the association of the another points type (*Peromyscus* species B) with a specific fixed location is randomized. In this case, the test does not address whether the *Peromyscus* species are themselves at random or clustered, but rather whether the association of type A points (*P. difficilis*) with type B points (*P. melanotis*) is what one would expect, given the locations of the points as fixed (Rosenberg and Anderson 2011). We assigned coordinates to each sampling stations considering 5 m distance between them, using the Cartesian coordinates of each sampling station where we captured mice as data input. Our null hypothesis was independence between *P. difficilis* and *P. melanotis* (Ripley 1976; Diggle 1983; Dale 1999); *i. e.,* we wanted to assess whether type A points (*P. difficilis*) were associated, or disassociated, with type B points (*P. melanotis*). In other words, Ripley’s K_12_(*d*) allowed us to assess spatial attraction or repulsion between the two species, among the stations of the plot. To evaluate the statistical significance of K_12_(*d*), we estimated the 95 % confidence interval (95 % CI), using a Monte Carlo procedure, based on 1000 stochastic relocation simulations of the sampling stations in the plot (Upton and Fingleton 1985; Zavala-Hurtado *et al*. 2000). When L(*d*) was positive and above the upper limit of the 95% CI, we inferred dissociation or repulsion between the two *Peromyscus* at the corresponding (*d*) scale; a significant negative deviation indicated a pattern of association or attraction between them (Dale 1999). If L(*d*) was within the 95 % CI for a given value of *d*, the null hypothesis of independence between the two contrasts could not be rejected (Dale 1999). The height of the L(*d*) function (peak height) indicates the intensity of the association or repulsion. We carried out a control for edge effect for the Ripley’s K analysis, rescaling the base counts overlapped with the study boundary (Rosenberg and Anderson 2011).

### Prediction of microhabitat elements affecting the use of space by each species

The next *s*tep was to assess which specific environmental variables best explained the frequency of each species within the complete grid at each season. For this, we performed Generalized Linear Models (GLMs), using JMP ® (ver. 9, SAS Institute Inc., 1989 - 2007, Cary, NC,). First, we calculated the mean for each one of the 23 variables in each microhabitat type, and then conducted a Principal Component Analysis (PCA) on the microhabitat means to obtain functions summarizing the most variance within each season. Because in both seasons the first three principal components (PC) summarized a good portion of the variance (38.61 for the dry season and 45.92 for rains), we used them to construct the GLMs. Since we found no multicollinearity between variables, we assessed different models using the PCs as explanatory variables to explain the frequency of individuals of each species for every season. Then, to visualize the fit of the model, the predicted values from the GLMs were plotted against the PC of the best model chosen. For each species, the response variable in the models was the capture frequency at each trapping station of the grid, and we assumed a Poisson distribution with a log link function.

Criteria for selecting the best models to explain the frequency of mice of each species in every season within the sampling stations included: a) model significance of *p <* 0.05; b) lowest Akaike Information Criterion (AIC), *i. e.,* a measure of goodness-of-fit penalized by the number of variables (Posada and Buckley 2004); c) Pearson goodness-of-fit (*p <* 0.05) and its deviation (Deviance; *p <* 0.05). If the models met all the criteria and showed very similar results, we selected the final model also taking into consideration the ecological and biological significance of the results.

## Results

### Frequency of captures

During the dry season, the total number of individuals was 111: 64 individuals for *P. difficilis* and 47 individuals for *P. melanotis*. During the rainy season, the total number of individuals increased to 168: 87 individuals for *P. difficilis* and 81 individuals for *P. melanotis* (Table A.2).

### Behavior of the 23 microhabitat variables between seasons

When considering the mean value over all stations, 16 out of the 23 environmental variables showed a significant difference between the two seasons (Table 2, Figure 3). Main observed changes involved variables related to herbaceous vegetation, according to plant coverage at different heights, and vegetation species richness (VR). As expected, overall, woody life forms were more stable between seasons (Table 2, Figure 3). Components of microhabitat showing no statistical change between seasons were frequency of herbaceous plants at 10 cm (HF11), frequency of woody plants at 35 cm (WF23) and 50 cm (WF24), as well as coverage of logs (CLO), rocks (RC), and bare soil (BS); therefore, space configuration given by these structural features remained stable in both seasons (Figure 3).

**Figure 3.**
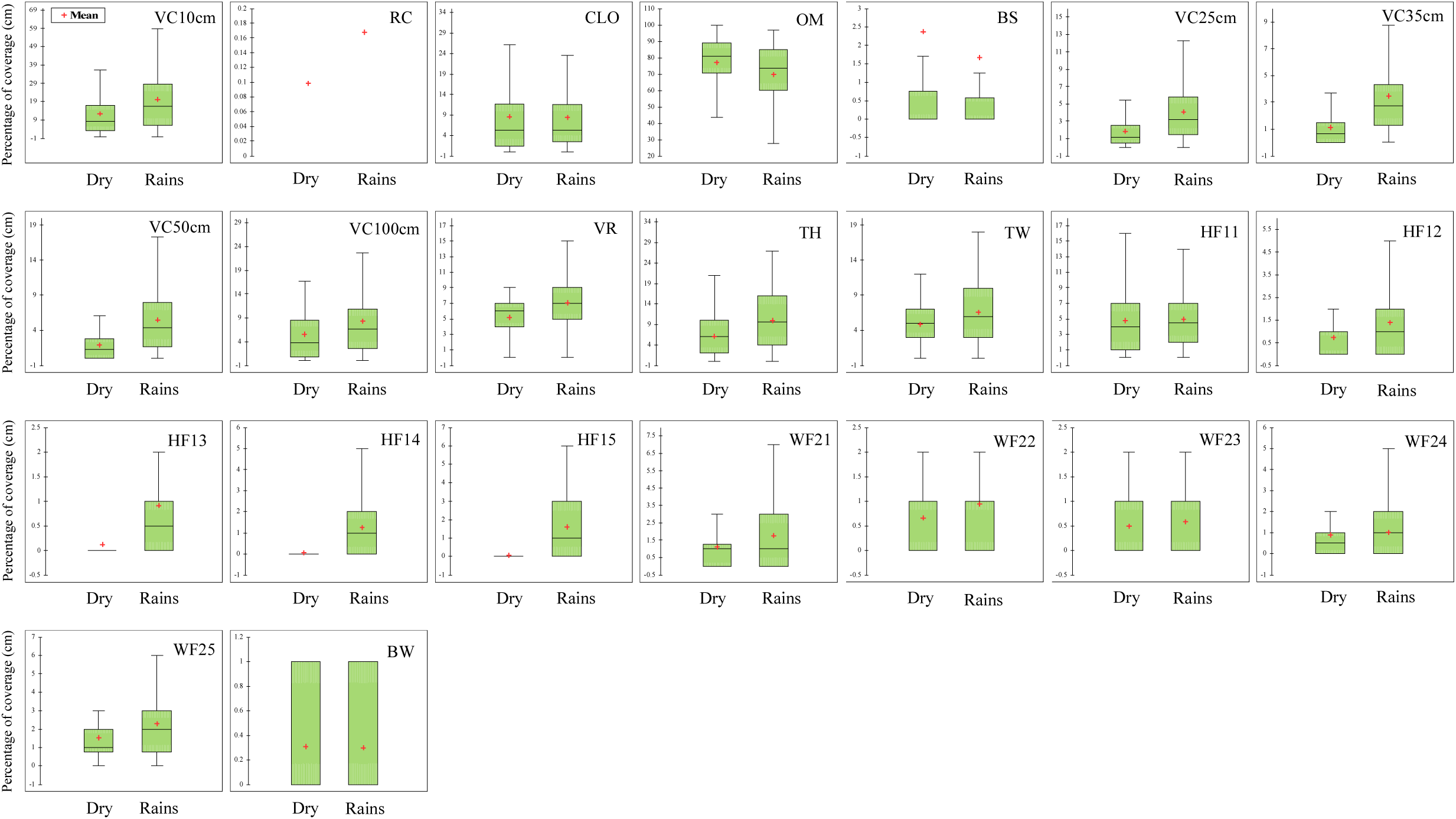
Seasonal heterogeneity between the dry and rainy seasons as evaluated using 23 environmental variables sampled in the study plot. Overall, vegetation variables increased during the rainy season, while organic matter (OM) decreased. Fixed elements, such as logs on the ground (CLO), showed no shifts between seasons. Variables with low values in the study plot (*e. g*., RC) are depicted with red crosses (mean value).

### Microhabitat heterogeneity

The three different microhabitat types (M1, M2, and M3) within the sampling plot in each season, as revealed by cluster analyses, revealed temporal habitat heterogeneity in the entire plot (Figure A.1). In addition, Fisher’s exact test of Contingency Tables Analysis revealed significant changes (*p* = 0.004) in the number of sampling stations assigned to each microhabitat type from the dry to the rainy seasons with main changes occurring within M1 and M2. That is, M1 included more sampling stations during the dry season *x*^2^ = 2.123, *p* < 0.05), while M2 covered the majority of sampling stations during the rainy season (*x*^2^ = 3.348, *p* < 0.05), as its area increased by 24 sampling stations from the dry to the rainy seasons (Figure 2, Figure A.1). Conversely, M3 remained quite stable (*x^2^* = 0.011), with almost the same number of sampling stations throughout the study (Figure 2).

Discriminant analyses validated heterogeneity of the three microhabitats (Figure 2) within and between seasons. The percentage of variance associated with each discriminant function was higher during the rainy season (dry season: F1 = 58.67 %, F2 = 42.32 %; rainy season: F1 = 72.581 %, F2 = 27.419 %, Figure A.1). Discriminant functions explained variation due to woody plants in the understory during the dry season, while herbaceous vegetation together with decayed matter and logs became more relevant during the rainy season (Table A.3). Wilk’s Lambda showed that at least the mean of one microhabitat differed statistically from the others either in the dry and rainy season (*p* = 0.0001 each, respectively, Figure A.1). These results corroborated spatial and temporal heterogeneity drawn from cluster analysis of individual station features in the grid plot. During the dry season and rainy season, 93.33 % and 94.17% of individual sampling stations, respectively, remained correctly classified (Figure 2).

### Description of microhabitats within the plot

*Microhabitat 1.* M1 (Figure 2, Figure A.1) was characterized by little to no vegetation cover at different heights (VC) and by a low frequency of herbaceous (HF) and woody (WF) plants. In contrast, organic matter on the ground (OM) was the most frequent component. This microhabitat was also distinctive for having extensive coverage of logs (CLO) on the ground and for being the microhabitat with the most conspicuous presence of burrows (BW).

*Microhabitat 2.* M2 (Figure 2, Figure A.1) was the largest area covered by herbaceous life forms at different heights (WF11-15); vegetation species richness (VR) and the total number of herbaceous plants (TH) also remained very high. Woody life forms at 25 cm (WF22) were present only at low frequencies, while WF24 (35 cm) and WF25 (100 cm) remained at higher frequencies. Coverages on the ground of small logs or dead wood (CLO), rocks (RC), and organic matter (OM) remained low during the study. Bare soil surface (BS) increased to higher amounts during the dry season and decreased during the rainy season. There were no burrows (BW) in this microhabitat.

*Microhabitat 3.* In M3 (Figure 2, Figure A.1), vegetation coverage showed high values at all height levels (VC25, VC35, VC50, and VC100), except for the first (VC10), while vegetation species richness (VR) averaged the highest, compared to other microhabitats. Total number of woody plants (TW) and frequency of woody plants at different heights (WF21 - 25) remained very high, while total number of herbaceous plants (TH) was low. Coverage of rocks (RC), logs (CLO), and bare soil surface (BS) were low during the dry season. Vegetation coverage at 10 cm (VC10) was also low during that season but increased in the rainy season. Conversely, OM was present in the dry season but decreased in the rainy season. There was a low number of burrows (BW).

### Association between the microhabitats and species

Kernel Graphs (Figure 4) indicated that the two species had a distinctive use of space, since they were differentially scattered along the plot. *Peromyscus difficilis* was more abundant during the dry season, occupying a large portion of the plot. *P. melanotis*, on the other hand, increased its distribution during the rains, when it seemed to displace *P. difficilis* into other sampling stations (Figure 4). Distribution rearrangements between seasons implied that one species occupied some stations more frequently than the other, and vice versa (Figure 4). Indeed, the two *x^2^* tests yielded statistically significant differences in abundance of each species among microhabitats and between seasons (Table A.2: dry season; R^2^ (U) = 0.09, *n* = 111, df = 2, Likelihood Ratio *x^2^* = 15.07, *p* = 0.00005; rainy season; R^2^ (U) = 0.02, *n* = 168, df = 2, Likelihood Ratio *x^2^* = 6.40, *p* = 0.0406).

**Figure 4.**
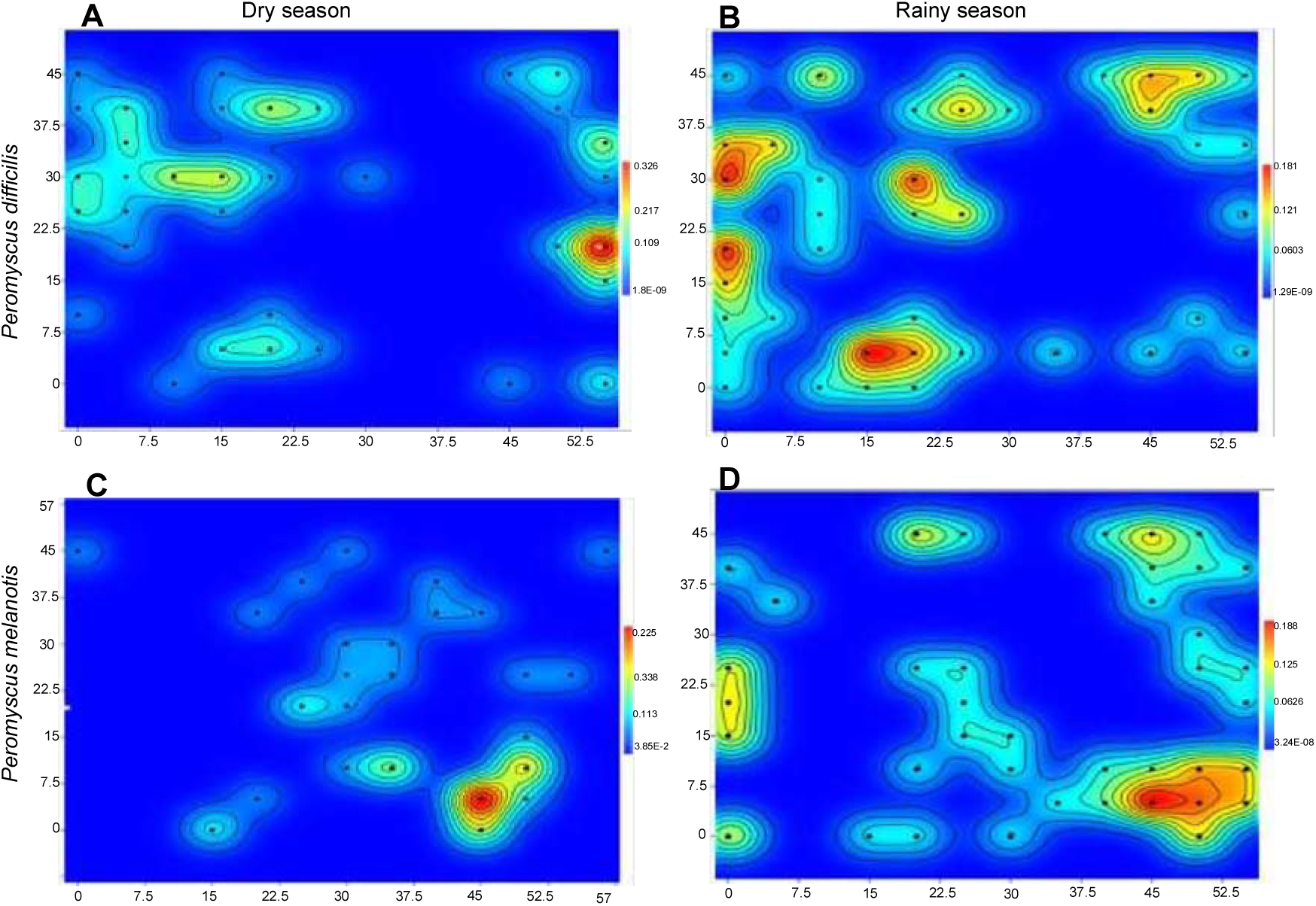
Spatial analysis with Kernel density points showing capture frequencies during the dry and rainy seasons for *P. difficilis* (A and B respectively) and *P. melanotis* (C and D, respectively). Dots depict actual capture points, while the color refers to the density of captures, with red color indicating the highest frequencies (hot spots). Numbers on each axis indicate actual distances (m).

During the dry season (Figure 5 and Table A.2), *Peromyscus difficilis* was highly associated with M2 (80 % of captured mice) and somewhat less with M1 (64 % of captured mice); conversely, *P. melanotis* was captured more frequently in M3 (66 % captured mice). Spatial use of microhabitats changed for both species in the rainy season (Figure 5 and Table A.2), also showing a microhabitat partition, though less conspicuous; *i. e*., in the Correspondence Analysis, 61 % of mice caught in M1 were *P. difficilis*. This species was also related to M3, while 60 % of mice caught of *P. melanotis* were captured in M2.

**Figure 5.**
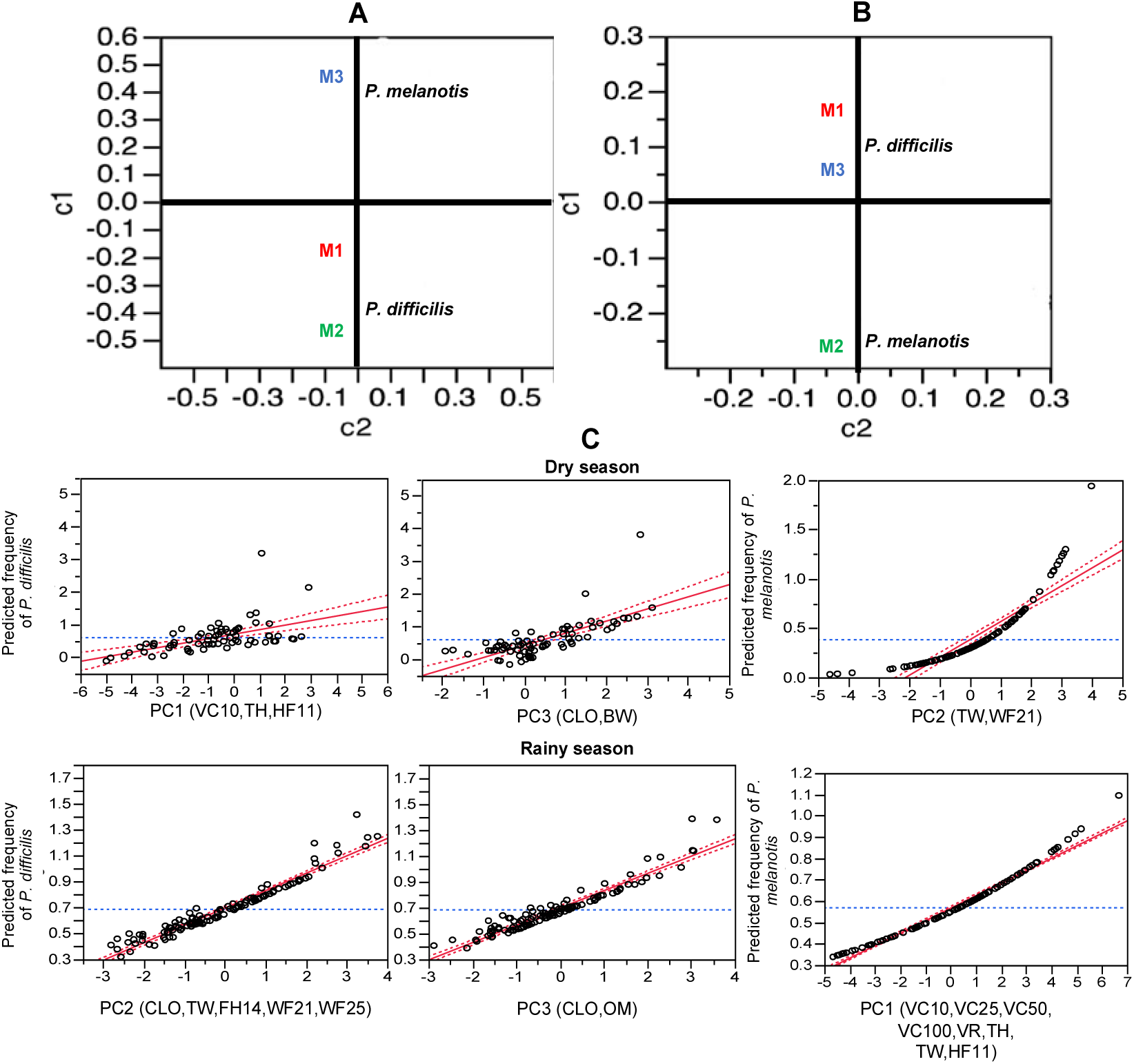
Correspondence analysis from contingency tables of capture frequency data for each deermouse (*P. difficilis* and *P. melanotis*) in each microhabitat (M1 - 3), during the dry (A) and rainy (B) seasons. Axes c1 and c2 indicate coordinates resulting from the ordination of each variable in the analysis. Plots for the GLMs models (C), showing predicted frequencies for each species, according to a determined principal component (model effect). All fitted PCs showed a positive relationship in the model, indicating that the probability of capture frequency of individuals of certain species, increased with the presence of the environmental variables within parenthesis (those having the highest loading scores).

### Spatial Patterns

The Nearest Neighbor Analysis revealed a significantly clustered pattern within the plot for each species in both seasons (Table 3). During the dry season, the mean distance between individuals of *P. difficilis* was more clustered than for individuals of *P. melanotis*. However, during the rainy season, the mean distance between individuals increased in *P. difficilis*, but decreased in *P. melanotis* (Table 3).

During the dry season, Ripley’s bivariate K showed a statistically significant pattern of repulsion between the *Peromyscus* species in almost all analyzed distances of the entire plot, except at 2 m where it showed an attraction pattern (Figure 6A). This repulsion pattern reversed in almost all distances during the rainy season (Figure 6B), with the two species becoming more associated, sharing microhabitats in almost all sampling stations. However, during the rainy season, statistically significant peaks of repulsion reappeared between the species at distances of five, ten, and 14 m (Figure 6B). These analyses also revealed the intensity of these patterns; *e.g.,* the likelihood of finding individuals of *P. difficilis* and *P. melanotis* together at the same capture station was very low during the dry season (Figure 6A), while this probability increased in the rainy season (Figure 6B).

**Figure 6.**
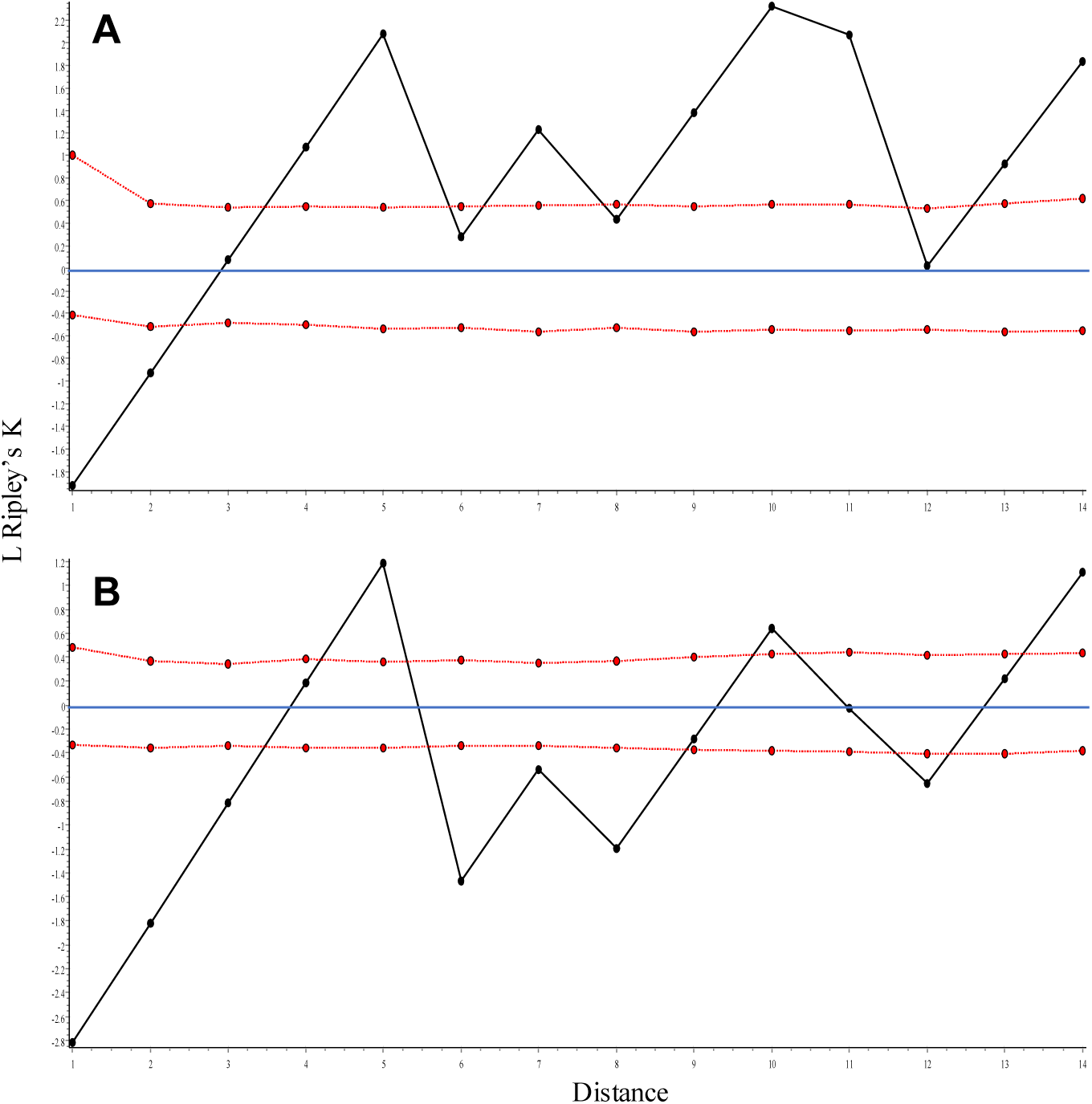
Ripley’s bivariate K Analysis for the interaction between the two *Peromyscus*. (A) shows a repulsion pattern between both species in all analyzed distances during the dry season. (B) Shows attraction from 6 - 12 m and repulsion at 5, 10, and 14 m, between the two species during the rainy season. The solid black line represents Ripley’s K bivariate pattern; red dashed lines depict a 95 % CI; solid blue line shows the null hypothesis. See Methods.

**Table 3.** Nearest Neighbor Analysis for the species of *Peromyscus*, showing the mean distance at what individuals of each species are dispersed among each other. In all instances, individuals of either species had a clustered point pattern. R - values indicate how the observed distribution deviates from random. Clustered points give *R* < 1, Poisson patterns give *R* ∼ 1, while overdispersed points give *R* > 1.

|  | Dry season |  | Rain season |  |
| --- | --- | --- | --- | --- |
|  | *P. difficilis* | *P. melanotis* | *P. difficilis* | *P. melanotis* |
| Points | 57 | 39 | 58 | 52 |
| Mean distance | 1.40 | 2.39 | 1.67 | 2.09 |
| $R$ | 0.42 | 0.77 | 0.55 | 0.64 |
| $P$ value | $1.1263 \times 10^{-16}$ | 0.006 | $8.6577 \times 10^{-11}$ | $8.7708 \times 10^{-7}$ |
| Point pattern | Clustered | Clustered | Clustered | Clustered |

### Effects of microhabitat structural components on distribution

According to the best Generalized Linear Model (AICc = 212.55, Table 5, Figure 5C), during the dry season PC1 and PC3 were the best predictors of capture frequency of *P. difficilis*. Environmental variables with the highest scores in PC1 (*p* = 0.0186) were the total number of herbaceous plants (TH), vegetation species richness (VR), and vegetation cover at 10 cm (VC10). In PC3 (*p* = 0.0001, Table 4, 5, Figure 5C), characterizing variables were the number of logs on the ground (CLO), and the presence of burrows (BW). These five variables were also the main elements characterizing M2 (PC1) and M1 (PC3), the two habitats with the highest capture frequencies of *P. difficils*. As for *P. melanotis* (Figure 5C), capture frequencies were positively related to PC2 (AICc = 204.47, *p* = 0.0001), and among the four variables with the highest PC scores were total number of woody plants (TW) and frequency of woody life forms at 10 cm height (WF21), the two main elements of M3 during the dry season (Table 5, Figure 5C).

**Table 4.** Principal components (PC1 - 3) used as effect model in the GLMs analysis for both seasons. Variable names as in Table 1.

| Variables | Dry season<br>(38.61 % of explained variance) |  |  | Rainy season<br>(45.92 % of explained variance) |  |  |
| --- | --- | --- | --- | --- | --- | --- |
|  | PC1 | PC2 | PC3 | PC1 | PC2 | PC3 |
| VC10 | 0.713 | -0.388 | -0.134 | 0.659 | -0.179 | -0.374 |
| RC | -0.145 | -0.055 | -0.110 | 0.014 | 0.062 | 0.034 |
| CLO | 0.030 | 0.118 | 0.828 | -0.153 | 0.411 | -0.583 |
| OM | -0.604 | 0.303 | -0.487 | -0.499 | -0.122 | 0.775 |
| BS | 0.130 | -0.187 | 0.112 | -0.028 | 0.154 | -0.328 |
| VC25 | 0.415 | 0.280 | 0.034 | 0.658 | -0.061 | 0.237 |
| VC32 | 0.287 | 0.184 | 0.052 | 0.580 | -0.003 | -0.126 |
| VC50 | 0.523 | 0.107 | 0.078 | 0.669 | -0.173 | -0.030 |
| VC100 | 0.335 | 0.408 | 0.052 | 0.656 | -0.053 | 0.107 |
| VR | 0.762 | 0.251 | -0.269 | 0.776 | 0.380 | 0.058 |
| TH | 0.837 | -0.430 | -0.131 | 0.874 | -0.319 | -0.076 |
| TW | 0.520 | 0.684 | -0.168 | 0.615 | 0.711 | 0.270 |
| HF11 | 0.784 | -0.394 | -0.214 | 0.678 | 0.007 | -0.228 |
| HF12 | 0.573 | -0.345 | 0.176 | 0.548 | -0.278 | 0.256 |
| HF13 | 0.247 | 0.073 | 0.113 | 0.463 | -0.260 | 0.067 |
| HF14 | 0.149 | -0.186 | -0.023 | 0.580 | -0.530 | -0.043 |
| HF14 | 0.080 | 0.117 | -0.236 | 0.481 | -0.267 | -0.019 |
| WF21 | 0.127 | 0.510 | -0.394 | 0.342 | 0.603 | 0.311 |
| WF22 | 0.015 | 0.064 | -0.105 | 0.266 | 0.212 | 0.416 |
| WF23 | 0.199 | 0.256 | -0.228 | 0.196 | 0.366 | -0.241 |
| WF24 | 0.424 | 0.359 | 0.195 | 0.337 | 0.165 | 0.159 |
| WF25 | 0.515 | 0.366 | 0.380 | 0.494 | 0.469 | 0.026 |
| BW | -0.105 | 0.249 | 0.557 | -0.170 | 0.332 | -0.218 |

**Table 5.**
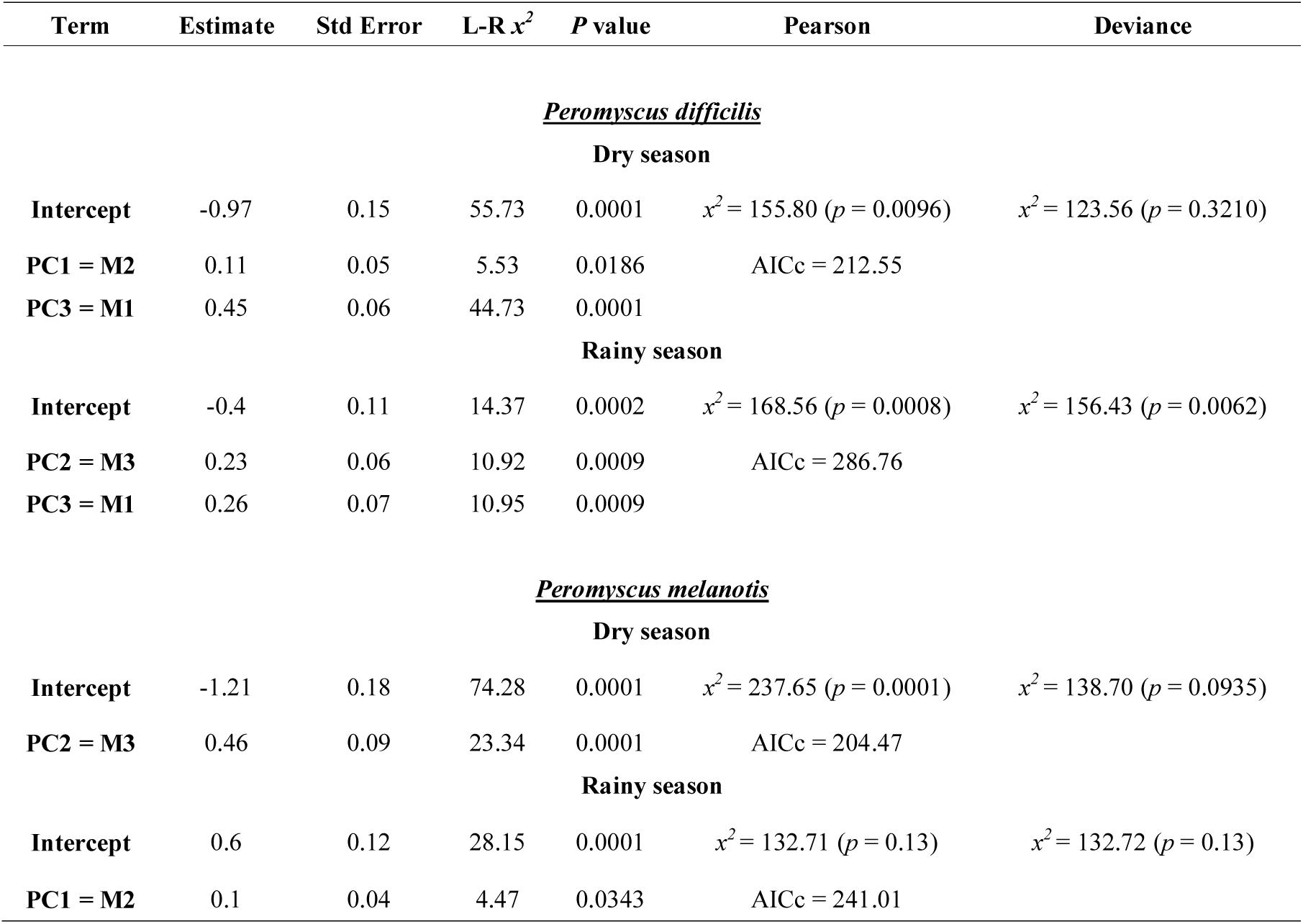
Generalized Linear Models constructed with the Principal Components (PC1 - 3, habitat resources) and frequencies of *P. difficils* and *P. melanotis* for each pluvial season. Resource partitions are observed in both species and seasons, since each PC relates to one microhabitat (M1 - M3); note how PC1 - 3 explain capture frequencies for each species within each season. Tests of Pearson and Deviance for fit goodness of are also shown.

During the rainy season, a similar pattern of microhabitat partition occurred between the two species. The best GLM model for *P. difficilis* in this season (Table 5) showed effects of PC2 and PC3. The most important environmental variables (AICc = 286.76, *p* = 0.0001, Table 4, 5) in PC2 were total number of woody plants (TW), herbaceous forms at 50 cm (HF14), woody forms at 10 and 100 cm (WF21, WF25, respectively), and number of logs on the ground (CLO). In PC3, the variables were number of logs on the ground (CLO) and organic matter (OM), which were associated with a higher capture frequency of *P. difficilis* in M1 (PC3) and M3 (PC2) during the rainy season. On the other hand, *P. melanotis* (Figure 5C) was only associated with PC1 (AICc = 241.01, *p* = 0.0343), and the main characterizing variables in this component were number of total herbaceous plants (TH), vegetation species richness (VR), and plant cover at almost all analyzed heights (VC10 - 100 cm, Table 4).

## Discussion

The results of our analyses show that as environmental conditions changed, the two syntopic species of *Peromyscus* switched and split space resources. Our findings on the microhabitat relationships between these two *Peromyscus*, dwelling in a patch of mid-latitude temperate forest, concur with the habitat heterogeneity coexistence hypothesis (Cramer and Willig 2002; Valladares *et al*. 2015). We found that both *Peromyscus* were sensitive to slight changes in microhabitat structure and that each one of the three microhabitat types provided different resources for each species. Our results also suggest that temporal differences in spatial structure, and the associated shifts in food and shelter availability, facilitate coexistence between these two syntopic *Peromyscus* (Pianka 1973; Schoener 1974). For instance, during the dry season, *P. difficilis* was most associatedwith M1 and M2, while *P. melanotis* was highly associated with M3. Conversely, in the rainy season, *P. difficilis* was more frequently captured in M1 and M3, while *P. melanotis* was highly associated with M2. Partition in space use has already been documented for coexisting species of *Peromyscus* within a community (*i. e.,* niche partitioning by *Peromyscus leucopus* and *P. maniculatus*, Kaufman and Kaufman 1989), and several studies have demonstrated variation in spatial resource use by congeneric species (Barry *et al*. 1990; Dooley and Dueser 1996; Kalcounis-Rüppell and Millar 2002).

In our study, *P. difficilis* remained highly associated with M1 in both seasons, but especially so during the rainy season. There was a very high number of fallen logs at M1, which represents small patches of microhabitat for small mammals with food sources, such as plant items and invertebrates (Bellows *et al*. 2001), as well as sources for refuge and shelter (Bowman *et al*. 2001). In this microhabitat, individuals of *P. difficilis* can also use the large fallen logs as pathways for quick and straight locomotion within the forest (Bellows *et al*. 2001; Dewalt *et al*. 2003). Indeed, fallen logs promote structural complexity of forests and may enhance positive interactions among species of small mammals (Bowman and Facelli 2013). GLM analyses showed that the main variables contributing to the abundance of *P. difficilis* during the rainy season were herbaceous vegetation (TH), vegetation species richness (VR), and the number of logs on the ground (CLO), among others. However, in the dry season, the main variables contributing to the abundance of *P. difficilis* were woody vegetation (TW), the number of logs on the ground, and organic matter (OM), among others. Bellows *et al*. (2001) found a similar result in a high latitude temperate forest (Virginia, USA), where distribution of a generalist small mammal was associated with the diameter of fallen logs, the frequency of shrubs, and degree of canopy closure. Association between rodents and fallen logs was also documented for *Nectomys squamipes*, which builds its nests inside decomposed fallen logs (Briani *et al*. 2004). This has also been recorded for other rodent species in several biomes with different vegetation types (*e. g*., *Rattus rattus*, *Nesomys audeberti* in Lehtonen *et al*. 2001; *Oligoryzomys nigripes* in Dalmagro and Vieira 2005). In contrast, *P. melanotis* was only related with one PC in each season. For instance, during the dry season, *P. melanotis* was associated with only PC2, a component representing mostly the total number of woody plants (TW) and small woody forms (WF21, 10 cm height). During the rainy season, the presence of *P. melanotis* was only related to PC1, which is characterized by an understory dominated by herbaceous life forms (TH). Higher rodent frequency in higher-density understory habitats has been interpreted as a strategy to avoid predation by aerial hunters (Grelle 2003; Dalmagro and Vieira 2005; Johnson 2007).

Other intrinsic factors, such as the breeding seasons, are likely to be related to coexistence of these two syntopic *Peromyscus*. For instance, the Nearest Neighbor Analysis revealed that in both seasons each species showed a distinctive, significant clustered pattern within the plot. Individuals of *P. difficilis* were more clustered together by mean distances than individuals of *P. melanotis*, particularly during the dry season. Conversely, during the rainy season, the mean distance between individuals increased in *P. difficilis* but decreased in *P. melanotis*. These shifting dispersion patterns are species-specific and relate with their respective breeding seasons (De-la-Cruz *et al*. 2019). The main breeding activity in *P. difficilis* occurs during the dry season, while it is during the rains for *P. melanotis*.

In addition, species-specific needs for different resource requirements to fulfill the breeding season, could affect the dispersion and coexistence between these two *Peromyscus* in the study area. For instance, coexistence between the two *Peromyscus* may be improved by temporal segregation of *P. melanotis*, compared to the resident behavior of *P. difficilis*. (species). Indeed, *P. melanotis* behaved as an opportunistic species throughout our study area since its capture frequencies increased when environmental conditions were more beneficial during the rainy season but remained at low levels during the harsher dry season. This smaller *Peromyscus* may migrate to areas with more availability of resources and better environmental conditions to survive during the dry season, coming back to the area during the rainy season to carry out the breeding season, when there is greater accessibility of food and spatial resources, and when competition with *P. difficilis* may decrease. On the other hand, as showed by Mohammadi (2010) in another small member of this genus, *Peromyscus leucopus*, opportunistic habitat use may be related to predation risk, which in our study is also supported for *P. melanotis* given its affinity for microhabitats that offer greater coverage against aerial predators. Further studies of the spatial segregation and predation influence on the opportunistic behavior of *P. melanotis* are needed.

Increase in number of captures for both species from the dry to the rainy seasons is due to resource availability, but they are also due to the type of resources. For instance, *P. difficilis* remained more abundant and always present at microhabitats with more stable elements in the study area throughout the study, while the increase in abundance of *P. melanotis* depended on denser plant cover during the rainy season. Since rains promote an increase of primary productivity, allowing more resource availability (mainly food) and enhancing microhabitat carrying capacity, such habitat changes facilitated coexistence between both *Peromyscus*, and with other small land mammals in the area (Castro-Campillo *et al*. 2008, 2012). Ripley’s bivariate K supported and shed light on this result, since the two species showed repulsion between them in almost all analyzed distances during the dry season; indeed, during the dry season when resources were scarce, incidents of both species using the same microhabitat were uncommon. Since the dry season is the primary breeding season for the medium-sized *P. difficilis*, it is likely that this species displays territorial behavior against its smaller-sized congeneric. Conversely, the intensity of this repulsion decreased substantially during the rainy season, with the two species showing an association pattern at some analyzed distances. This supports the idea that increased resource availability during the rainy season allows the two species to share the habitat. Moreover, our results also indicate that spatial dispersion patterns are seasonally variable in both *Peromyscus* (Brown and Zeng 1989; Cramer and Willig 2002). In this sense, it is important to point out that in both seasons Ripley’s bivariate K indicated an attraction pattern between the two species within 2 and 3 m, which is more likely an artefact of our sample design than a biological result. This is because if an individual of *P. difficilis* was captured during the first of our two sampling nights, followed by capture of one individual of *P. melanotis* in that same station the next day, the analysis counted the pattern as an association because they were trapped in the same station at a very short distance.

Also, partition of space by these syntopic deermice must be facilitated by their respective locomotive habits-semiarboreal in *P. difficilis* and cursorial in *P. melanotis*. The long tail of *P. difficilis* enables it to rush and climb along shrubs or trees (Fernandez *et al*. 2010), thus increasing its preference for habitats with these and other fixed (*e. g.,* fallen logs and twigs) elements where it can escape from predators. In fact, adult coat color changes in *P. difficilis* between seasons, becoming more similar to ground litter, suggesting that in this resident deermouse, phenotype plasticity in color is a cryptic response elicited by predation risk. In contrast, smaller body size, together with a shorter tail and narrower soles (Álvarez-Castañeda 2005), should enable the cursorial *P. melanotis* to occupy such zones as M2 and M3, where predators cannot easily spot it through dense vegetation cover, so it can scape very quickly. Indeed, coverage by high shrubs provides both protection from predators and food sources, since seeds may be concentrated under shrub canopies (Thompson 1982; Mohammadi 2010). In fact, rodents usually avoid foraging in unsheltered microhabitats and forest edges where they are more likely to be spotted by avian (Kotler *et al*. 1991) and other vertebrate predators (Morris and Davidson 2000; Mohammadi 2010).

Finally, several aspects of our methods for sampling the environmental features of microhabitat, as well as the quantitative analyses used here, especially Ripley’s bivariate K, make it particularly informative in studies of spatial dynamics of dispersion in small rodents at a microhabitat scale. Our explicitly designed methodological approach, focused on small-sized rodents with low vagility, provided us with necessary information about the ability of these two *Peromyscus* species to split resources in the complex understory of a mid-latitude temperate forest. Our initial aim to reconstruct microhabitat structure also successfully provided us with clues as to how to eventually manage perturbed ecosystems for conservation purposes, such as those at the edge of a megalopolis, whose continuous growth produces fragmentation of natural microhabitats. We think that our methodology will also be helpful in other scenarios for understanding the ecology of small rodents, such as behavioral dynamics, activity patterns, and reproductive systems at intra- and/or inter-population levels.

## Acknowledgments

We thank C. Peralta-Juarez and J. Patiño-Ortega for their friendly fieldwork assistance and logistic support. The staff at UAMI mammal scientific collection kindly provided all the facilities for specimen curation and housing. Authorities in charge of DLNP (CORENA, SEMARNAT), as well as the forest rangers and administration staff at the National Park gave us necessary information, security, and logistics to complete our studies in the field. This paper contains partial information from the Masters in Biology thesis (DCBS, UAM-I) of IMDA; IMDA received financial support through a fellowship (283799, CVU 479479) from the Consejo Nacional de Ciencia y Tecnología (CONACyT). This study was supported by annual grants (143.***.46 IB-DCBS-UAMI, 2013-2017) to AACC.

