## supplemental material figures for "Spatiotemporal microhabitat heterogeneity and dispersion patterns of two small mammals in a temperate forest"

**Appendix. Table A.1.** Clustered sampling stations in the study plot, within each of three microhabitats (M1 - 3), according to 23 environmental variables, in two pluvial seasons. See figure 2.

| M1 | M2 | M3 |
|---|---|---|
| **Dry season** | | |
| M1 (N = 68) | M2 (N = 28) | M3 (N = 24) |
| A1,B5,A2,B2,A3,B3,C4,B4,D4,C5,C3C9,C10,A4,B1,A6,J4,K8,A10,G6,D1,H6,H7,B8,D9,H2,H3,I3,B6,H8,E8,G9,J7,J3,J10,A7,B10,D10,G1,G3,H1,I1,I8,I7,K9,J8,C8,G7,G10,I4,L8,E10,L9,I10,A9,H9,F7,J6,A5,H10,K7,A8,C1,C2,E7,D2,E2,F9 | B7,C6,C7,L5,K1,L6,L4,D8,K4,K5,K6,E6,I2,L3,I6,J5,L7,F4,L10,F3,I5,J9,F6,H5,F8,G4,G8,D7 | B9,E4,F5,D6,I9,L2,D3,D5,E3,H4,K3,E1,F1,E5,E9,F2,K2,G2,F10,J1,J2,G5,K10,L1 |
| **Rainy season** | | |
| M1 (N = 46) | M2 (N = 51) | M3 (N = 23) |
| A1,A2,B5,C1,A8,D10,B10,A3,B1,B4,C4,B3,C3,B8,H9,B9,E8,K8,C9,C10,A4,A6,C5,A5,A9,A10,E10,A7,J8,L1,G5,I4,C2,D9,F7,J7,G6,G7,E4,E9,I7,D7,H3,B2,L8,E7 | B6,H6,H7,C6,C7,D6,J5,I5,C8,K3,G4,F8,I6,H8,K7,F4,F6,I1,G8,L3,G10,H10,D4,J4,L6,J2,K9,I2,I3,I8,J6,L7,J3,J10,K1,H2,K2,B7,L5,J9,L4,E3,F1,G9,I10,F3,K4,D8,I9,K5,K6 | D1,D3,D5,E5,E6,L9,L10,F2,G1,H1,H5,D2,K10,F10,F5,F9,G2,J1,E1,H4,L2,E2,G3 |

**Table A.2.** Contingency Analysis with $x^2$ tests, showing captures of mice in each microhabitat (M1 - 3), during the respective dry and rainy seasons. See figure 5.

| | | ***Peromyscus difficilis*** | ***Peromyscus melanotis*** | |
|---|---|---|---|---|
| | | **Dry Season** | | **Total** |
| | **Count** | 32 | 18 | |
| **M1** | **Row %** | 64 | 36 | 50 |
| | $x^2$ | 0.349 | 0.47 | |
| | **Count** | 20 | 5 | |
| **M2** | **Row %** | 80 | 20 | 25 |
| | $x^2$ | 2.164 | 2.94 | |
| | **Count** | 12 | 24 | |
| **M3** | **Row %** | 33.33 | 66.67 | 36 |
| | $x^2$ | 3.69 | 5.03 | |
| | **Total** | 64 | 47 | 111 |
| | | **Rainy Season** | | **Total** |
| | **Count** | 37 | 23 | 60 |
| **M1** | **Row %** | 61.67 | 38.33 | |
| | $x^2$ | 1.13 | 1.21 | |
| | **Count** | 24 | 37 | 61 |
| **M2** | **Row %** | 39.34 | 60.66 | |
| | $x^2$ | 1.82 | 1.95 | |
| | **Count** | 26 | 21 | 47 |
| **M3** | **Row %** | 55.32 | 44.68 | |
| | $x^2$ | 0.11 | 0.12 | |
| | **Total** | 87 | 81 | 168 |

**Table A.3.** Factor (F1 - 3) loadings for 23 environmental variables (VAR) in the Discriminant Analysis for three microhabitats during the dry (Dry) and rainy (Rains) seasons, respectively. Highlighted numbers in red point out variables with highest scores; *i. e.*, those with the most discriminant effect in each season for the *a priori* groups from the Clustering Analysis.

| Dry | | | Rains | | | |
|---|---|---|---|---|---|---|
| **VAR** | **F1** | **F2** | **VAR** | **F1** | **F2** | **F3** |
| **VR** | 0.698 | -0.165 | **TH** | 0.884 | -0.062 | -0.115 |
| **TW** | 0.615 | 0.271 | **CV10** | 0.714 | -0.066 | -0.067 |
| **HF11** | 0.563 | -0.671 | **HF14** | 0.689 | 0.207 | -0.071 |
| **WF21** | 0.562 | 0.408 | **HF11** | 0.664 | -0.212 | -0.085 |
| **TH** | 0.500 | -0.718 | **HF12** | 0.567 | 0.056 | -0.064 |
| **CV10** | 0.459 | -0.578 | **VR** | 0.563 | -0.633 | -0.031 |
| **CV50** | 0.417 | -0.322 | **CV50** | 0.522 | -0.068 | -0.049 |
| **HF15** | 0.406 | 0.363 | **CV25** | 0.501 | -0.167 | -0.034 |
| **WF24** | 0.379 | -0.034 | **HF15** | 0.481 | 0.004 | -0.087 |
| **CV100** | 0.314 | 0.202 | **HF13** | 0.443 | 0.070 | 0.010 |
| **WF25** | 0.280 | -0.163 | **CV100** | 0.398 | -0.143 | -0.054 |
| **CV35** | 0.221 | -0.041 | **TW** | 0.294 | -0.872 | -0.002 |
| **HF12** | 0.214 | -0.554 | **WF4** | 0.242 | -0.369 | 0.003 |
| **CV25** | 0.212 | -0.034 | **CV35** | 0.226 | -0.204 | -0.028 |
| **HF13** | 0.102 | -0.185 | **WF25** | 0.204 | -0.617 | -0.008 |
| **BS** | 0.100 | -0.159 | **RC** | 0.203 | -0.002 | -0.021 |
| **WF23** | 0.091 | 0.091 | **WF22** | 0.200 | -0.302 | -0.014 |
| **HF14** | -0.084 | -0.113 | **WF21** | 0.115 | -0.737 | 0.011 |
| **BW** | -0.125 | 0.206 | **WF23** | 0.090 | -0.109 | -0.009 |
| **RC** | -0.130 | 0.115 | **BS** | -0.064 | -0.170 | 0.011 |
| **CLO** | -0.133 | 0.039 | **BW** | -0.261 | -0.224 | 0.053 |
| **WF22** | -0.150 | 0.025 | **CLO** | -0.315 | -0.059 | 0.036 |
| **OM** | -0.298 | 0.475 | **OM** | -0.452 | 0.151 | 0.044 |

**Table A.4.** The different Generalized Linear Models (GLMs) tested for *P. difficilis* and *P. melanotis* in the dry and rains are observed. Different models were used to build the GLMs.

| *P. difficilis* | | | | | | |
|---|---|---|---|---|---|---|
| **Dry season** | | | | | | |
| Term | Estimate | Std Error | L-R $x^2$ | *P* value | Pearson | Deviance |
| **Intercept** | -1.04 | 0.16 | 59.66 | 0.001 | $x^2$ = 152.26 (p = 0.0135) | $x^2$ = 116.19 ($p$ = 0.4775) |
| **PC1** | 0.10 | 0.05 | 3.52 | 0.0605 | AICc = 207.31 | |
| **PC2** | 0.25 | 0.095 | 7.37 | 0.0066 | | |
| **PC3** | 0.45 | 0.066 | 43.03 | 0.0001 | | |
| **Intercept** | -0.80 | 0.14 | 42.27 | 0.0001 | $x^2$ = 201.88 ($p$ = 0.0001) | $x^2$ = 159.22 ($p$ = 0.0058) |
| **PC1** | 0.22 | 0.06 | 13.79 | 0.0002 | AICc = 248.21 | |
| **PC2** | 0.23 | 0.07 | 9.07 | 0.0026 | | |
| **Intercept** | -0.97 | 0.15 | 55.73 | 0.0001 | $x^2$ = 155.80 ($p$ = 0.0096) | $x^2$ = 123.56 ($p$ = 0.3210) |
| **PC1** | 0.11 | 0.05 | 5.53 | 0.0186 | AICc = 212.55 | |
| **PC3** | 0.45 | 0.06 | 44.73 | 0.0001 | | |
| **Intercept** | -1.04 | 0.16 | 59.36 | 0.0001 | $x^2$ = 160.08 ($p$ = 0.0050) | $x^2$ = 119.71 ($p$ = 0.4130) |
| **PC2** | 0.29 | 0.09 | 9.38 | 0.0022 | AICc = 208.70 | |
| **PC3** | 0.49 | 0.06 | 53.30 | 0.0001 | | |
| **Rainy season** | | | | | | |
| **Term** | Estimate | Std Error | L-R $x^2$ | *P* value | Pearson | Deviance |
| **Intercept** | -0.41 | 0.11 | 15.19 | 0.0001 | $x^2$ = 167.50 ($p$ = 0.0013) | $x^2$ = 161.21 ($p$ = 0.0035) |
| **PC1** | -0.06 | 0.04 | 2.19 | 0.1385 | AICc = 286.71 | |
| **PC2** | 0.18 | 0.06 | 7.81 | 0.0052 | | |
| **PC3** | 0.16 | 0.06 | 6.67 | 0.0098 | | |
| **Intercept** | -0.40 | 0.11 | 14.01 | 0.0002 | $x^2$ = 179.79 ($p$ = 0.0002) | $x^2$ = 167.89 ($p$ = 0.0014) |
| **PC1** | -0.07 | 0.04 | 2.80 | 0.0938 | AICc = 291.24 | |
| **PC2** | 0.23 | 0.06 | 11.40 | 0.0007 | | |
| **Intercept** | -0.38 | 0.11 | 13.38 | 0.0003 | $x^2$ = 181.01 ($p$ = 0.0001) | $x^2$ = 169.03 ($p$ = 0.0012) |
| **PC1** | -0.06 | 0.04 | 2.24 | 0.1344 | AICc = 292.28 | |
| **PC3** | 0.22 | 0.06 | 10.26 | 0.0014 | | |
| **Intercept** | -0.40 | 0.11 | 14.37 | 0.0002 | $x^2$= 168.56 ($p$ = 0.0008) | $x^2$= 156.43 ($p$ = 0.0062) |
| **PC2** | 0.23 | 0.06 | 10.92 | 0.0009 | AICc = 286.76 | |
| **PC3** | 0.26 | 0.07 | 10.95 | 0.0009 | | |

| *P. melanotis* | | | | | | |
|---|---|---|---|---|---|---|
| **Dry season** | | | | | | |
| **Term** | **Estimate** | **Std Error** | **L-R $x^2$** | ***P* value** | **Pearson** | **Deviance** |
| **Intercept** | -1.22 | 0.18 | 74.97 | 0.0001 | $x^2$ = 220.26 (*p* = 0.0001) | $x^2$ = 137.33 (*p* = 0.0860) |
| **PC1** | 0.06 | 0.08 | 0.59 | 0.4408 | AICc = 207.33 | |
| **PC2** | 0.44 | 0.09 | 22.2 | 0.0001 | | |
| **PC3** | -0.10 | 0.10 | 0.97 | 0.3242 | | |
| **Intercept** | -1.21 | 0.18 | 74.44 | 0.0001 | $x^2$ = 230.79 (*p* = 0.0001) | $x^2$ = 138.30 (*p* = 0.0871) |
| **PC1** | 0.05 | 0.08 | 0.40 | 0.5235 | AICc = 206.17 | |
| **PC2** | 0.45 | 0.09 | 22.69 | 0.0001 | | |
| **Intercept** | -0.98 | 0.15 | 61.29 | 0.0001 | $x^2$ = 295.18 (*p* = 0.0001) | $x^2$ =159.61 (*p* = 0.0054) |
| **PC1** | 0.07 | 0.06 | 1.02 | 0.3114 | AICc = 227.48 | |
| **PC3** | -0.12 | 0.10 | 1.38 | 0.2397 | | |
| **Intercept** | -1.21 | 0.18 | 74.66 | 0.0001 | $x^2$ = 228.35 (*p* = 0.0001) | $x^2$ = 137.92 (*p* = 0.0001) |
| **PC2** | 0.45 | 0.09 | 22.71 | 0.0001 | AICc = 205.79 | |
| **PC3** | -0.09 | 0.11 | 0.78 | 0.3757 | | |
| **Intercept** | -1.21 | 0.18 | 74.28 | 0.0001 | $x^2$ = 237.65 (*p* = 0.0001) | $x^2$ = 138.70 (*p* = 0.0935) |
| **PC2** | 0.46 | 0.09 | 23.43 | 0.0001 | AICc = 204.47 | |
| **Rainy season** | | | | | | |
| **Intercept** | -0.61 | 0.12 | 28.95 | 0.0001 | $x^2$ = 136.70 (*p* = 0.0919) | $x^2$ = 132.90 (*p* = 0.1350) |
| **PC1** | 0.10 | 0.04 | 4.95 | 0.0260 | AICc = 248.04 | |
| **PC2** | -0.01 | 0.07 | 0.02 | 0.8803 | | |
| **PC3** | -0.14 | 0.09 | 2.66 | 0.1028 | | |
| **Intercept** | -0.58 | 0.12 | 27.75 | 0.0001 | $x^2$ = 134.80 (*p* = 0.1245) | $x^2$ = 135.56 (*p* = 0.1155) |
| **PC1** | 0.10 | 0.04 | 4.93 | 0.0262 | AICc = 248.56 | |
| **PC2** | -0.001 | 0.07 | 0.000 | 0.9887 | | |
| **Intercept** | -0.60 | 0.12 | 28.94 | 0.0001 | $x^2$ = 136.29 (*p* = 0.1072) | $x^2$ = 132.92 (*p* = 0.1491) |
| **PC1** | 0.10 | 0.04 | 4.94 | 0.0261 | AICc = 245.92 | |
| **PC3** | -0.14 | 0.09 | 2.63 | 0.1043 | | |
| **Intercept** | -0.62 | 0.13 | 29.51 | 0.0001 | $x^2$ = 136.30 (*p* = 0.0854) | $x^2$ = 132.22 (*p* = 0.1299) |
| **PC2** | -0.05 | 0.08 | 0.34 | 0.5596 | AICc = 242.61 | |
| **PC3** | -0.23 | 0.10 | 4.95 | 0.0260 | | |
| **Intercept** | -0.60 | 0.12 | 28.15 | 0.0001 | $x^2$ = 132.71 (*p* = 0.1375) | $x^2$ = 132.72 (*p* = 01372) |
| **PC1** | 0.10 | 0.04 | 4.47 | 0.0343 | AICc = 241.01 | |
| **Intercept** | -0.62 | 0.12 | 29.19 | 0.0001 | $x^2$ = 135.73 (*p* = 0.1017) | $x^2$ = 132.56 (*p* = 0.1394) |
| **PC3** | -0.22 | 0.10 | 4.64 | 0.0312 | AICc = 240.84 | |

**Figure A.1.** Classification of the three types of microhabitats (M1 = red, M2 = green, M3 = blue), in the dry (a) and rainy (b) seasons, respectively. The upper figure shows the behavior of the 23 variables within each influence zone for both seasons. The X-axis displays the three microhabitats resulted from cluster analysis and in Y-axis are plotting the 23 standardized variables, and error bars are indicated in each bar. This plot shows that all variables are standardized with mean = 0 and standard deviation = 1. Therefore, tends above the mean, indicate what that specific variable is above the mean in that types of microhabitats. While below 0 shows the opposite way. Below cluster analysis is observed the classification by Discriminant analysis of each microhabitat.

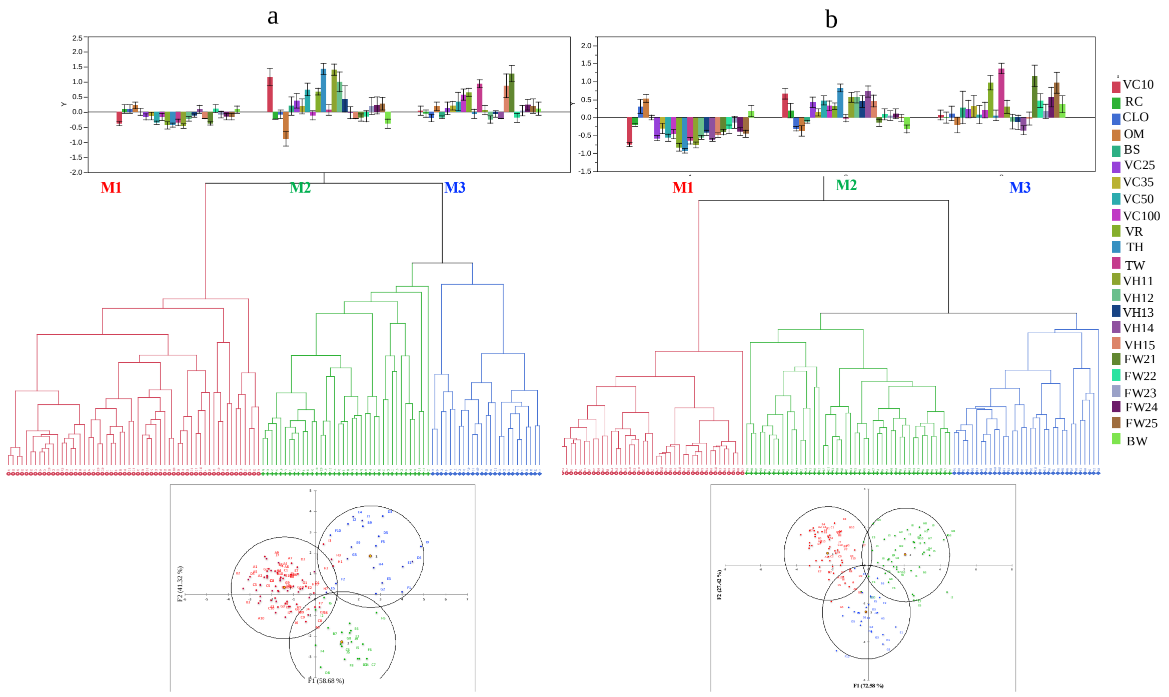

**Figure A.2.** Variance accumulation of the Principal Components Analysis of each new function for the dry season (a) and the rainy season (b). The first three principal components in both seasons explain the higher variance in the models (38.61 for the dry season and 45.92 for rains). Therefore, the first three PCs were taken to perform the GLIMs. Figures c and d show the Biplot graphs of the variables associated with the first two CPs (CP1 and CP2) and the spatial ordering of the sampling stations in these CPs is observed.

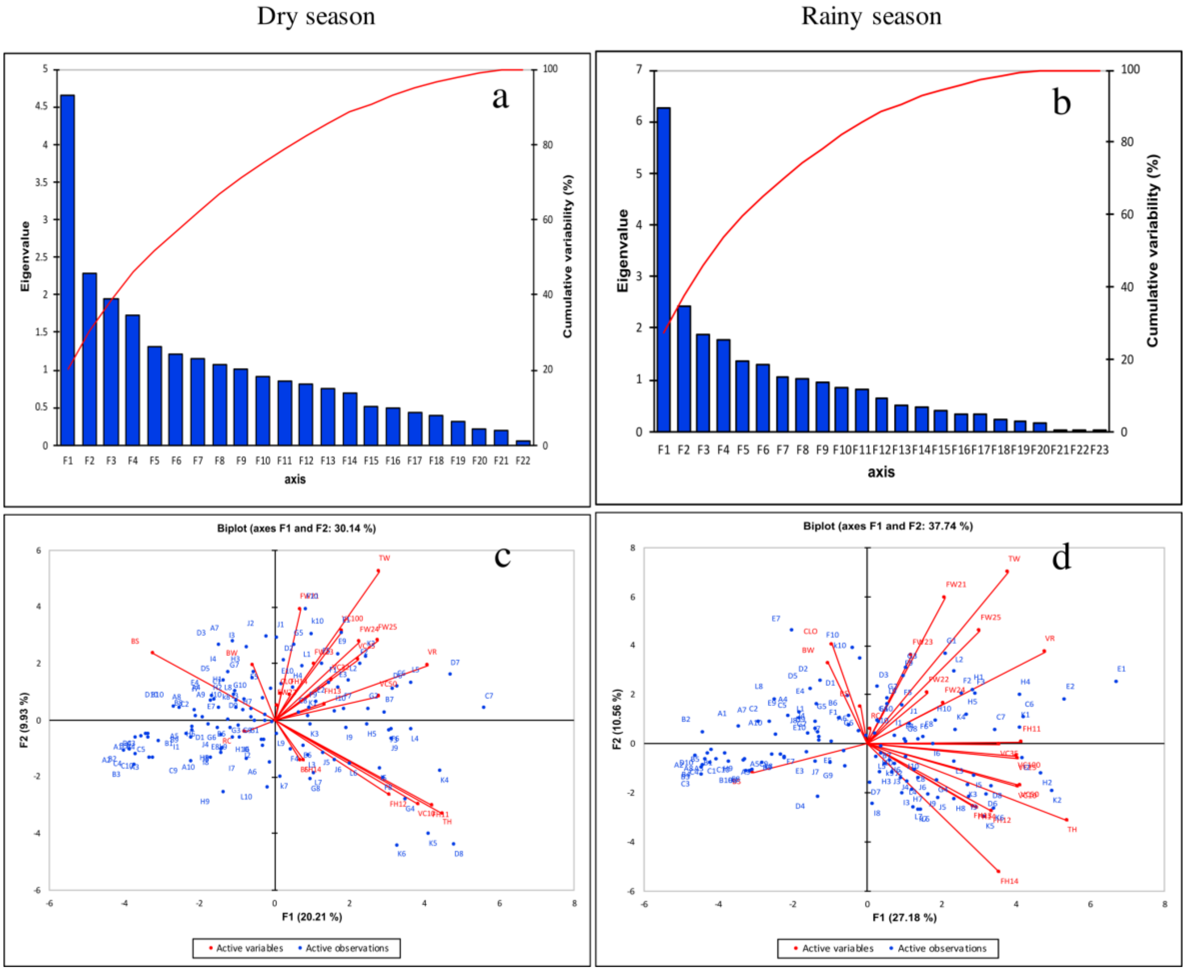
